# Large transgene arrays cause aberrant transcription and synthetic molting defects with *myrf-1* mutation

**DOI:** 10.64898/2026.04.30.721998

**Authors:** Iskra Katic, Panagiotis Papasaikas, Dimos Gaidatzis, Helge Großhans

## Abstract

Multicopy transgene arrays remain widely used in *C. elegans* research. It is usually assumed that they behave neutrally, not impacting the phenotype under investigation. Here, we reveal that a previously reported heterochronic extra-molt phenotype associated with *myrf-1(mg412)* depends on the presence of the integrated molting reporter *mgIs49*. Nanopore long-read sequencing shows that *mgIs49* is a massive 8.8-Mb insertion - around 50% of the size of its host chromosome - which disrupts the *prmt-9* gene. Both *mgIs49* and another array, *maIs105*, cause dysregulation of the transcriptome and accumulation of reads mapping to the promoter sequences used as components of the array. We identify additional arrays exceeding 4 Mb and show that variable molting defects occur across different transgenic lines when combined with *myrf-1(mg412)*, implicating array size or composition in the synthetic phenotype. Our results underscore the necessity of replacing multicopy reporters in developmental studies with single-copy insertions or endogenous tagging whenever possible.

## Introduction

Transgenes are a mainstay of modern biological research, permitting perturbation as well as read-out of gene expression or cellular states. In *C. elegans*, genome editing (Arribere et al. 2014; Dickinson et al. 2015; Paix et al. 2015; Dokshin et al. 2018; Ghanta and Mello 2020) or transposon-based single copy transgene insertion (Frøkjaer-Jensen et al. 2008; Frøkjær-Jensen et al. 2012) have become the methods of choice for many of these applications, as they permit targeted manipulation and physiological expression levels.

This contrasts with earlier approaches using transgene arrays consisting of multiple, concatenated copies of the transgene of interest (Mello et al. 1991). Following microinjection of DNA into the *C. elegans* germline, these arrays can be maintained extrachromosomally, but random segregation causes mosaic inheritance. This can be prevented by random integration of the array into of the genome, following X- or gamma ray (Mello and Fire 1995) or UV light/TMP treatment (Evans 2006) to induce DNA double-strand breaks. Certain problematic features of these integrated arrays are well understood: a high transgene copy number, which may also include truncated or otherwise abnormal copies, render them less suitable for mechanistic studies on gene expression and the repetitive nature makes them susceptible to heterochromatinization and small RNA-mediated silencing, especially in the germline (Kelly et al. 1997; Kim et al. 2005; Ashe et al. 2012).

Yet, many multicopy integrated transgene arrays are still routinely used as gene expression and cell fate reporters because decades of research using them have established them as convenient tools and even de facto gold standards. For instance, in our field of research, *C. elegans* developmental timing, we and others have found multicopy integrated arrays immensely valuable to monitor molting, larval vs adult gene expression states, cell numbers or fates, and numerous publications have shown their value for this purpose. The underlying assumption for these experiments is that the transgenic array does not, or not substantially, alter the phenotype under investigation. Here, we show that this premise does not hold when we study the role of *myrf-1* in molting.

*myrf-1* encodes a transcription factor that plays important roles in developmental timing control through the heterochronic pathway and in molting (Frand et al. 2005; Meng et al., 2017; Xia et al., 2021; Xu et al., 2024). Whereas strong hypomorph and null mutants fail to proceed through the normal four larval molts, previous work identified a single point mutation, *myrf-1(mg412)*, that reactivated a larval molt reporter array, *mgIs49[mlt-10p::gfp::pest; ttx-3::gfp]*, in adult animals, causing an aberrant molt and animal death (Frand et al. 2005).

Here, we show that the phenotype arises synthetically only in the presence of both *myrf-1(mg412)* and the *mgIs49* array. Although mapping reveals that *mgIs49* disrupts the *prmt-9* gene on Chromosome IV, we exclude this as the cause for the phenotype. Instead, various arrays cause synthetic phenotypes with *myrf-1(mg412)*, to variable extents. We find that integrated array sizes are consistently huge, ranging from 4 MB to 11 MB for each of eight examples that we mapped, accounting in the case of *mgIs49* for > 50% of the 17.49 MB of the wild-type chromosome IV (and > 5% of the entire genome size). Moreover, for two arrays examined in greater detail, *mgIs49* and the unrelated *maIs105[col-19p::gfp]* (Abbott et al., 2005; Ilbay et al. 2021), we observe substantial dysregulation of gene expression and accumulation of reads covering the promoter sequences utilized in the arrays even in the presence of wild-type *myrf-1*.

Collectively, the *myrf-1* synthetic phenotype appears best explained by generic features of the arrays such as large size or concatenation presumably at the core of the aberrant promoter transcription. Although we are reassured by the fact that the phenotype is not explained by the array alone but does additionally require the *myrf-1* mutation, our findings strongly suggest that multicopy arrays have outlived their usefulness even as tissue or cell type markers and should be replaced whenever possible by single copy transgenes or endogenously tagged genes.

## Results

### *The* mgIs49 *transgene array is required for the* myrf-1(mg412) *mutant phenotype*

Recently, we identified *myrf-1* as a candidate component of an oscillator, or clock, that times rhythmic development, especially molting, in *C. elegans* (Meeuse et al. 2023). Specifically, we found that RNAi-mediated depletion of this rhythmically transcribed gene caused animals to arrest development or die molting, consistent with earlier work showing that a *myrf-1(tm2707)* putative null mutation caused early larval lethality (Russel et al. 2011). (Russel et al. 2011) had additionally conducted a screen for molting regulators based on inappropriate activation of an integrated multicopy transgene, *mgIs49[mlt-10p::gfp::pest; ttx-3::gfp]*. (We will refer to this transgene as *mlt-10p::gfp::pest* in the remainder of this article.) They identified animals carrying a mutation in *myrf-1* (named *pqn-47* at the time), *mg412*, that caused reactivation of this larval molt reporter in adults, initiation of a new molt, and ultimately death, when animals became trapped in their cuticle. In other words, *myrf-1(mg412)* appeared to exhibit a retarded heterochronic phenotype, where adult animals failed to exit from the molting cycle.

In seeking to study the molecular details of the function of MYRF-1, we outcrossed the *myrf-1(mg412)*; *mgIs49* animals to wild-type N2. Surprisingly, we found animals that were homozygous for the mutation but showed no evidence of cuticle entrapment and premature death when cultured on plates (Figure 1a). This suggested that an unlinked mutation was partly or fully responsible for this phenotype. Notably, these healthy animals lacked the *mgIs49* array, suggesting that the array itself, or features linked to it, could contribute to the effect.

**Figure 1.**
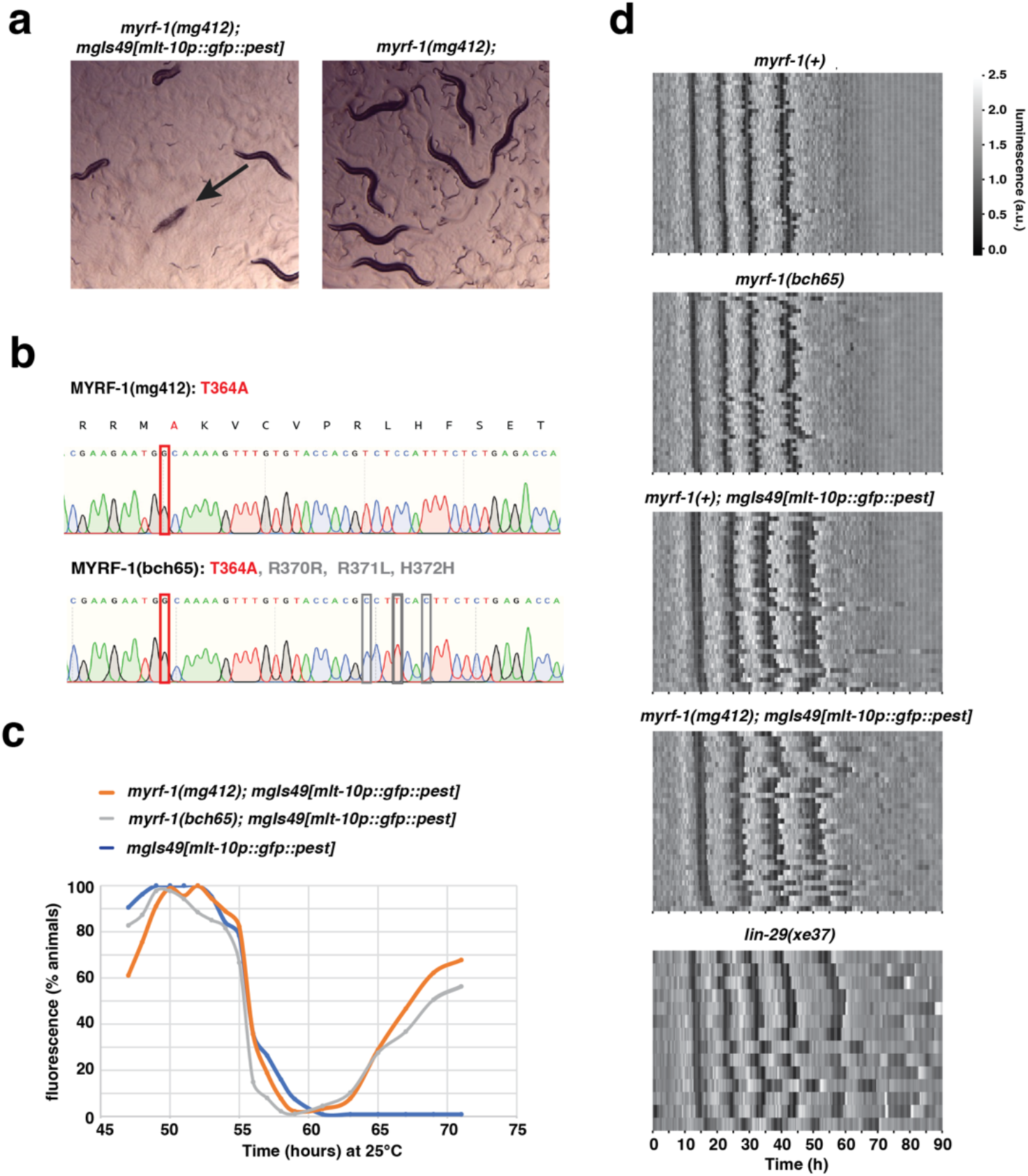
Adult lethality in *myrf-1(mg412)* animals is dependent on the presence of the *mgIs49[mlt-10p::gfp::pest]* array. **a**.Micrograph showing characteristic carcasses observed when growing *myrf-1(mg412)*; *mgIs49[mlt-10p::gfp::pest]* animals but not observed for *myrf-1(mg412)* animals. **b**.Chromatogram of relevant *myrf-1(mg412)* and *myrf-1(bch65)* sequences confirms CRISPR-mediated regeneration of the T364A missense change of *mg412* in *bch65* (*red boxes*). *myrf-1(bc65)* additionally contains three silent mutations (*black boxes*). **c**.Reactivation of the *mgIs49[mlt-10p::gfp::pest]* reporter in the presence of *myrf-1* alleles at adulthood. Percentage of animals (n>87 for each genotype) showing GFP fluorescence from *mgIs49[mlt-10p::gfp::pest]* at indicated times. The animals were grown at 25°C and scored for GFP expression from 47 to 72 hours using a stereomicroscope. Table 1b shows survival outcomes for these animals at 96 hours. **d**.Heatmap showing trend-corrected luminescence for the indicated strains (n>13), expressing luciferase from the *eft-3* promoter. Each line represents one animal and traces are sorted by entry into first molt. Darker color represents low luminescence and is associated with molts. *lin-29(xe37)* animals dying at the juvenile-to-adult transition were censored.

To investigate whether *myrf-1(mg412)* was at all required for the phenotype seen in *myrf-1(mg412); mgIs49* animals, we reverted the *myrf-1* mutation in that strain back to the wild-type coding sequence. This suppressed lethality in adults grown at 25°C (Table 1). Together these observations indicated that the *myrf-1* mutation was required but not sufficient for the phenotype.

**Table 1:** *myrf-1(mg412)* acts synergistically with *mgIs49[mlt-10p::gfp::pest]* to cause an adult lethality phenotype.

| Relevant genotype* | % death at 96 hours, 25°C (n) |
| --- | --- |
| *myrf-1(mg412); mgls49[mlt-10p::gfp::pest]* | 98 (165) |
| *mgls49[mlt-10p::gfp::pest]* | 14 (262) |
| *myrf-1(bch71[WT revertant of mg412]); mgls49[mlt-10p::gfp::pest]* | 21 (279) |
| *myrf-1(bch72[WT revertant of mg412]); mgls49[mlt-10p::gfp::pest]* | 32 (218) |
| *myrf-1(bch73[WT revertant of mg412]); mgls49[mlt-10p::gfp::pest]* | 22 (219) |

b
| Genotype | % death at 96 hours, 25°C (n) |
| --- | --- |
| *mgls49[mlt-10p::gfp::pest]* | 4 (106) |
| *myrf-1(mg412); mgls49[mlt-10p::gfp::pest]* | 98 (90) |
| *myrf-1(bch65); mgls49[mlt-10p::gfp::pest]* | 84 (87) |
| *myrf-1(bch65)* # | 0 (226) |

\* All lines also contain *xeSi311[eft-3p::luc::gfp::unc-54 3'UTR, unc-119(+)] V*

We tested this hypothesis further by regenerating the *mg412* mutation in an otherwise wild-type (non-transgenic) animal (Figure 1b) and subsequently crossing the new *myrf-1(bch65)* allele into *mgIs49*. The results were fully consistent with our hypothesis, since *myrf-1(bch65)* alone did not exhibit adult lethality, whereas *myrf-1(bch65); mgIs49[mlt-10p::gfp::pest]* did, along with adult *mlt-10p::gfp::pest* reactivation (Table 1; Figure 1c). These results also support the phenotype arising from a combination of *myrf-1* mutation and presence of the transgene array.

We also note that the heterochronic extra molt phenotype reported for *myrf-1(mg412); mgIs49[mlt-10::gfp::pest]* differs from that of a canonical retarded heterochronic mutation, *lin-29* null (Azzi et al. 2020): When we assessed molting in a high-throughput luciferase assay (Olmedo et al. 2015; Meeuse et al. 2020), *lin-29(xe37)* control animals showed the expected extra molts, whereas neither *myrf-1(bch65)* nor *myrf-1(mg412); mgIs49[mlt-10::gfp::pest]* animals did (Figure 1d). However, *mgIs49* caused developmental delays irrespective of *myrf-1* mutational status (Supplemental Figure 1).

**Supplementary Figure 1.**
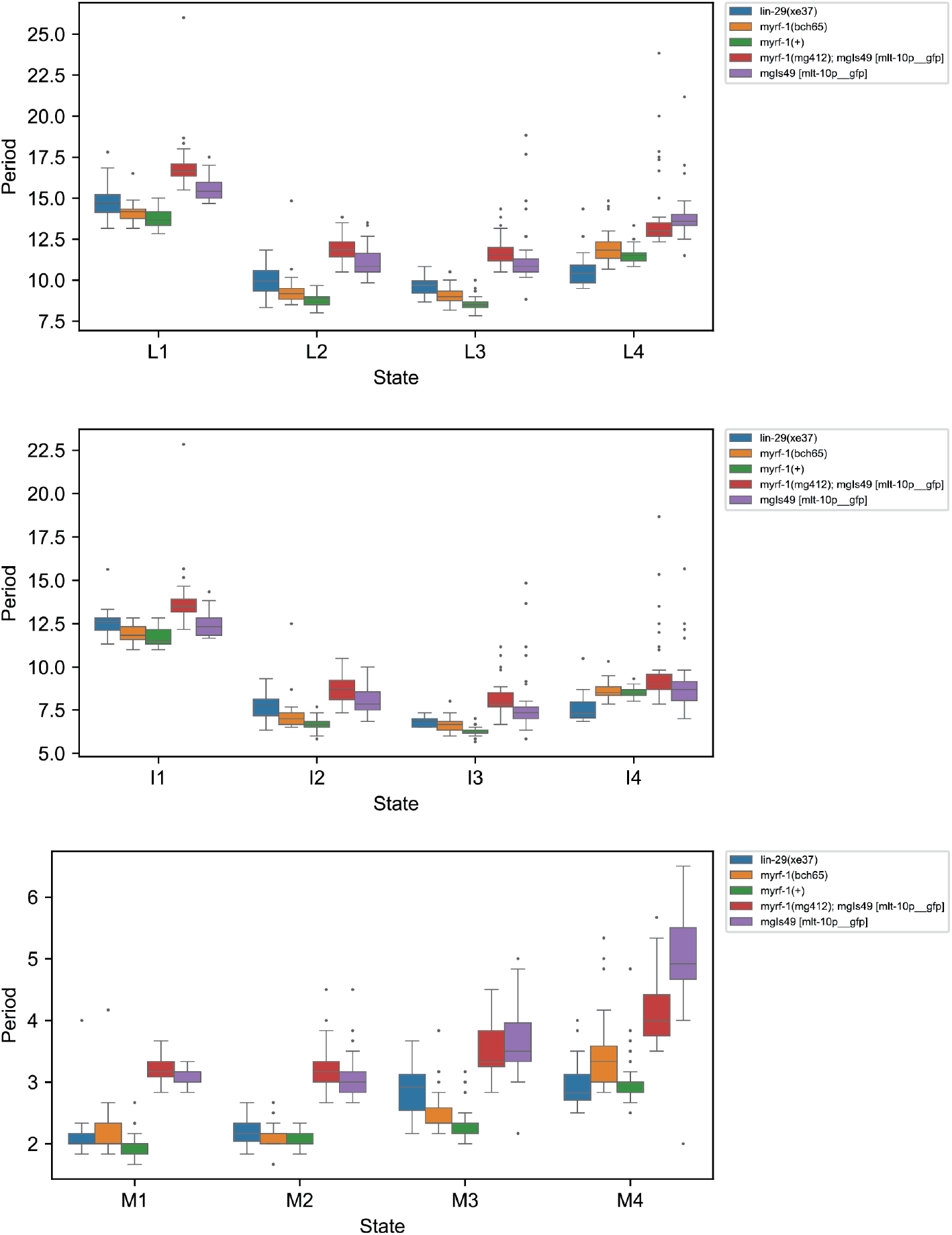
The *mgIs49[mlt-10p::gfp::pest; ttx-3p::gfp]* array causes developmental delay. Boxplots show larval stage length, intermolt length and molt length for animals of the indicated genotypes (n>13), expressing luciferase from the *eft-3* promoter. *lin-29(xe37)* animals dying at the juvenile-to-adult transition were censored.

### A mlt-10p::gfp MosSCI reporter does not synergize with myrf-1(mg412)

To investigate whether *myrf-1* mutation could reactivate molting gene expression independently of the *mgIs49* array, we used MosSCI to integrate a single-copy *mlt-10p::pest::gfp::h2b* transgene in a targeted fashion into a landing site on chromosome III (Frøkjær-Jensen et al. 2012). We characterized two transgenic lines, neither of which caused lethality, irrespective of whether this was assessed in a *myrf-1* wild-type or *myrf-1(mg412)* or *myrf-1(bch65)* mutant background (Table 2). At the same time, there was no reactivation of the transgene in any of the adults (Figure 2). We conclude that not only the lethality phenotype but also reactivation of molting gene expression requires *mgIs49* along with *myrf-1(bch65)* or *myrf-1(mg412)* mutations.

**Table 2:** Neither a *mlt-10p::gfp* MosSCI reporter nor a *prmt-9* deletion synergize with *myrf-1(mg412)* to cause an adult lethality phenotype.

| Genotype | % death at 96 hours, 25°C (n) |
| --- | --- |
| *bchSi134[mlt-10p::pest::gfp::h2b]* | 0 (188) |
| *myrf-1(mg412); bchSi134[mlt-10p::pest::gfp::h2b]* | 1 (159) |
| *bchSi134[mlt-10p::pest::gfp::h2b]* | 0 (155) |
| *myrf-1(bch65); bchSi134[mlt-10p::pest::gfp::h2b]* | 0 (158) |
| *bchSi135[mlt-10p::pest::gfp::h2b]* | 0 (224) |
| *myrf-1(mg412); bchSi135[mlt-10p::pest::gfp::h2b]* | 1 (255) |
| *bchSi135[mlt-10p::pest::gfp::h2b]* | 0 (282) |
| *myrf-1(bch65); bchSi135[mlt-10p::pest::gfp::h2b]* | 0 (288) |

b
| Relevant genotype* | % death at 96 hours, 25°C (n) |
| --- | --- |
| *myrf-1(bch65)* | 0 (226) |
| *myrf-1(bch65); prmt-9(bch79)* | 3 (144) |
| *myrf-1(bch65); prmt-9(bch80)* | 2 (149) |
| *myrf-1(bch65); prmt-9(bch81)* | 4 (143) |

\* All lines also contain *xeSi311[eft-3p::luc::gfp::unc-54 3'UTR, unc-119(+)] V*

**Figure 2.**
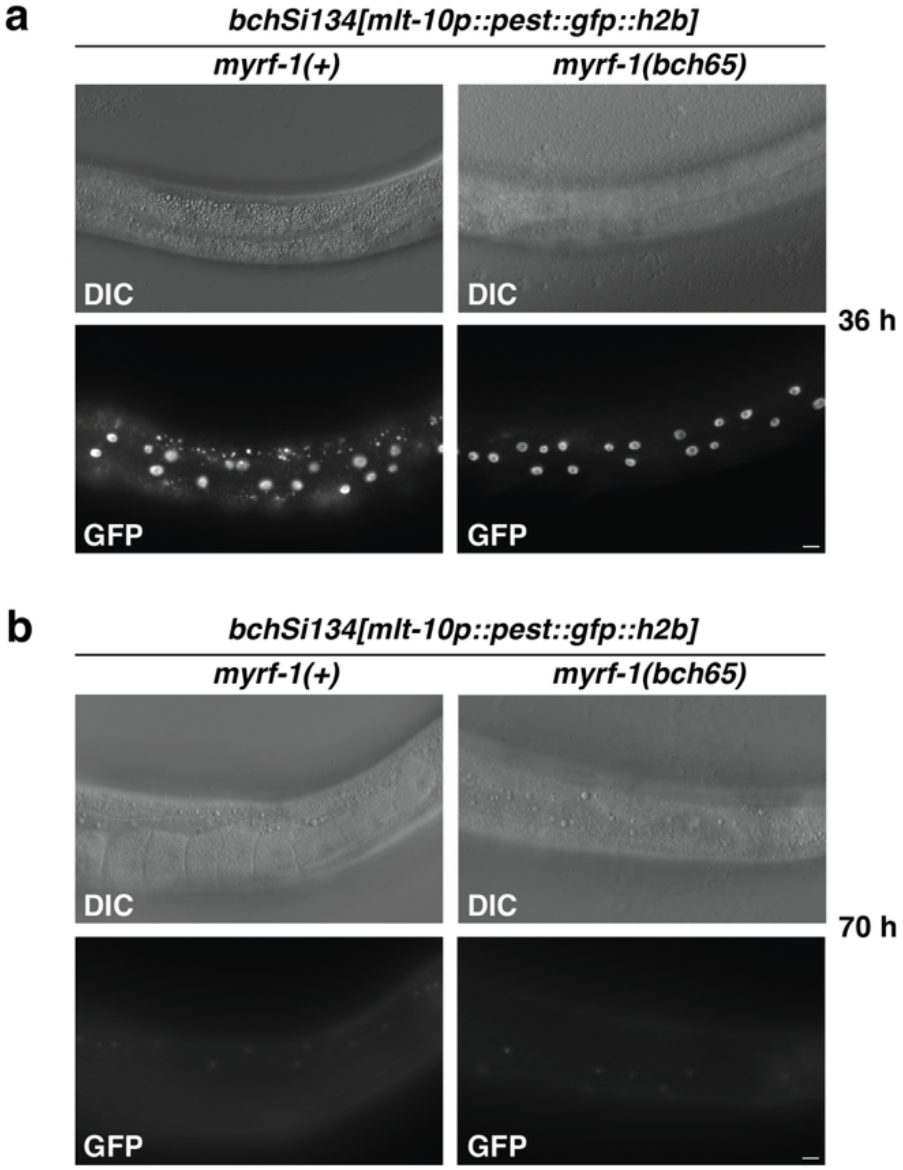
A single-copy reporter of *mlt-10* does not get reactivated in adulthood by *myrf-1(bch65)*. **a**., **b**. *bchSi134[mlt-10p::pest::gfp::h2b]* III reporter expression in WT and *myrf-1(bch65)* larvae, respectively, grown at 25°C for 36 hours (a) or 70 h (b) after plating starved L1s. Images were acquired using the same exposure times for GFP. Scale bars: 10 µm.

### mgIs49 *is a large integrated array disrupting* prmt-9

To further characterize *mgIs49*, we used ONT long-read sequencing to map the insertion site(s) and estimate the total size of the integrated array (see Materials and Methods). This analysis revealed integration of a large array into a protein-coding gene (*prmt-9)* (Figure 3). The array’s size was estimated at ~8.8 Mb, or around 50% of the chromosome’s wild-type size (Supplementary Table 1).

**Figure 3.**
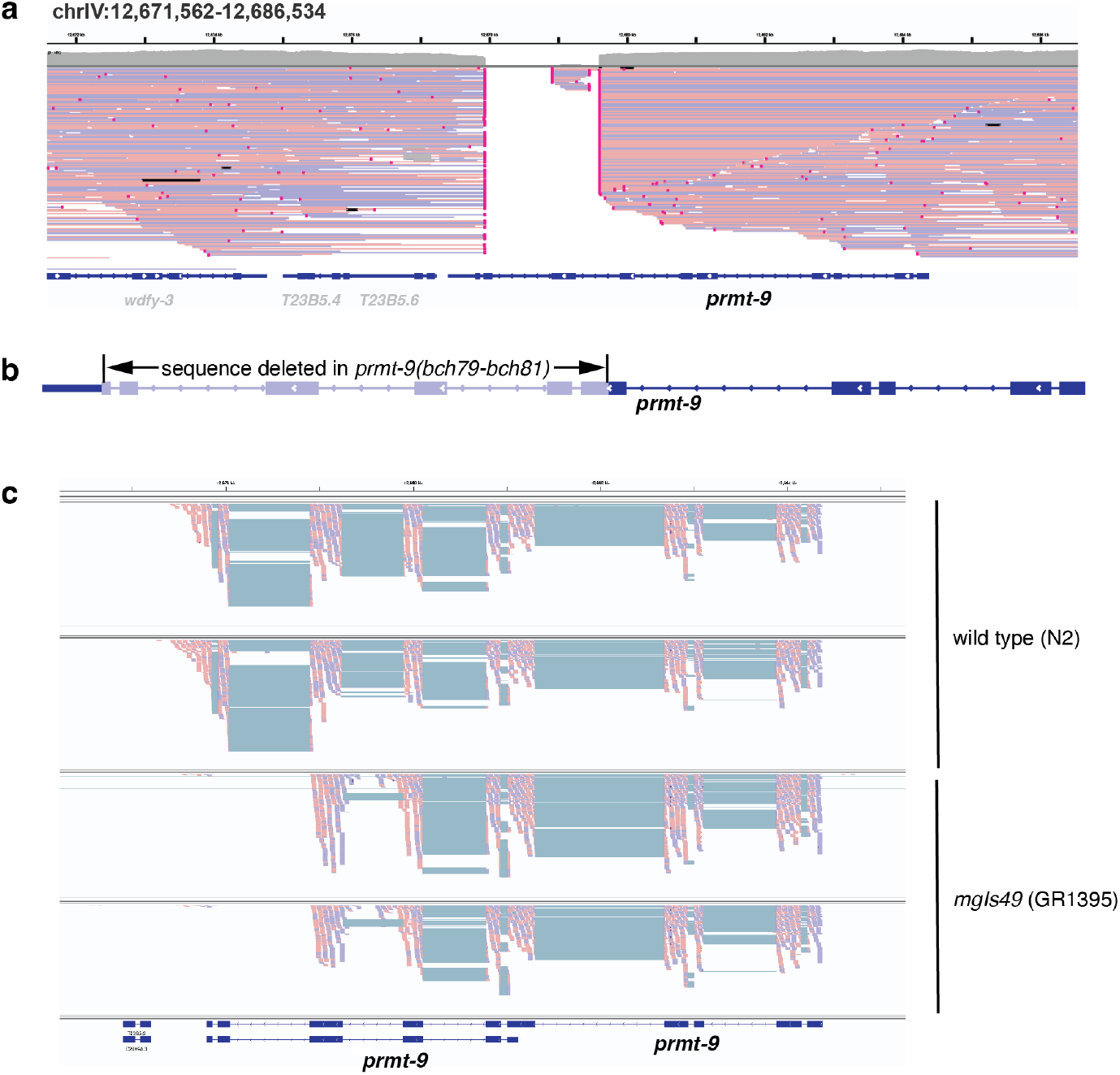
*mgIs49[mlt-10p::gfp::pest]* insertion disrupts the *prmt-9* coding sequence on chromosome IV. **a**.IGV genome browser screenshot of the identified strain-specific integration in GR1395. Red and blue lines show ONT long reads in the + and – orientation respectively and their primary mapping to the genome; genes are shown on the bottom of the plot. The characteristic breakpoint alignment structures indicate soft-clipped reads flanking the insertion site of the transgene array. In many cases the clipped sequences can be mapped to components of the array and no other strain analyzed in this study shows a breakpoint pattern on this locus (data not shown). **b**.Schematic of the CRISPR/Cas9 editing of the *prmt-9* coding sequence, showing its longer isoform. The alleles *bch79, bch80 and bch81* contain exons 1-4, a partial exon 5, and a partial exon 10 of *prmt-9*. **c**.mRNA-seq reads mapping to the *prmt-9* sequence in GR1395 *mgIs49[mlt-10p::gfp::pest]* and N2 (two replicates each). Reads corresponding to exons 9 and 10 of the longer *prmt-9* isoform are absent in GR1395.

To test whether *prmt-9* inactivation caused the observed genetic interaction, we disrupted it by genome editing in wild-type animals. This failed to result in adult lethality both on its own and in combination with *myrf-1(bch65)* (Table 2), suggesting that *prmt-9* mutation is not, or not alone, the cause of the observed phenotypes arising in presence of *mgIs49*.

### *Other transgene arrays synergize with* myrf-1(mg412) *to variable extents*

It seemed possible that the synthetic interactions between *myrf-1* mutation and *mgIs49* were specific to this combination. Alternatively, transgenic arrays might more broadly cause phenotypes when combined with *myrf-1*. To distinguish between these possibilities, we crossed two other integrated arrays, expressing different transgenes, into *myrf-1(bch65)*. Both arrays also caused synthetic adult lethality with *myrf-1(bch65)*, although the effect was more moderate with *maIs105[col-19::gfp]* (Abbott et al. 2005) than with *feIs5[sur-5p::luciferase::gfp + rol-6(su1006)]* (Lagido et al. 2008) (Table 3).

**Table 3:**
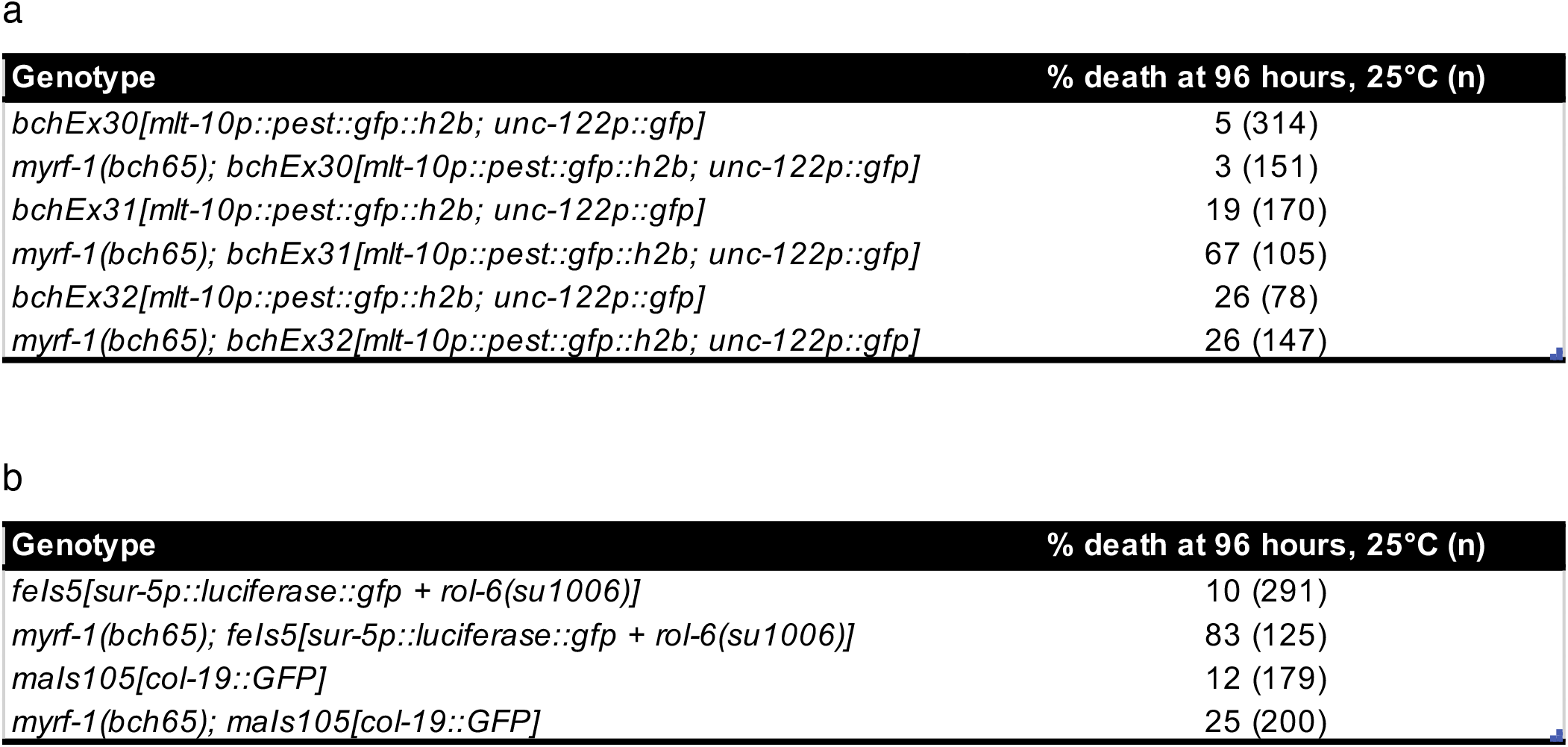
Extrachromosomal and integrated arrays cause adult lethality phenotype in the presence of *myrf-1(mg412)* mutation.

We wondered whether the synthetic phenotypes required array integration and therefore created extrachromosomal lines bearing *mlt-10p::pest::gfp::h2b*. Similar to *mgIs49*, each of the three extrachromosomal lines showed some degree of lethality at 96 hours even in a *myrf-1* wild-type background. In one of the three lines, this phenotype was enhanced if *myrf-1(bch65)* was present.

Taken together, we conclude that several but not all transgene arrays cause synthetic phenotypes with *myrf-1* mutation. This implicates a more generic feature than the specific molecular composition or integration site of *mgIs49* in the process, possibly including a large physical size and a repetitive nature.

### Integrated multicopy transgenes are megabases in size

To get a broader view of the physical size range of transgene arrays, we applied the same ONT long-read sequencing and array characterization pipeline to six additional transgene strains used in the community. The results of this survey suggest that the size that we had determined for *mgIs49* was in no way unusual: the estimated insertion sizes for each of the arrays range from ~4 to ~11 Mb, accounting for a significant fraction of the entire genome in those animals, and are on the same order of magnitude as individual chromosomes, which range from 14 to 21 Mb.

The long-read sequencing data also allowed us to map candidate array integration sites (Supplementary Table 2). We selected *maIs105[col-19p::gfp]* for further validation, using SNP mapping (Davis et al. 2005) to test a putative insertion on the right arm of chromosome V (Figure 4). We crossed *maIs105* animals with CB4856 Hawaiian isolate males and selected, based on their GFP-expression status, F_2_ animals either homozygous for the array or devoid of it. The Hawaiian SNP snp_Y17D7B [3] (Davis et al. 2005) showed a strong segregation bias with GFP: While only 1/34 animals homozygous for the array segregated the Hawaiian SNP, 36/36 F_2_ animals lacking the array were homozygous for it. Hence, these data confirm that *maIs105* is tightly linked to this SNP in the *Y17D7B*.*3* gene (whose interpolated genetic position is V: 17.75). Of note, two additional candidate insertion sites were detected on chromosome II, along with multiple candidate integration sites in several other strains, consistent with the idea that array integration events can produce complex genome aberrations (Supplementary Table 2).

**Figure 4.**
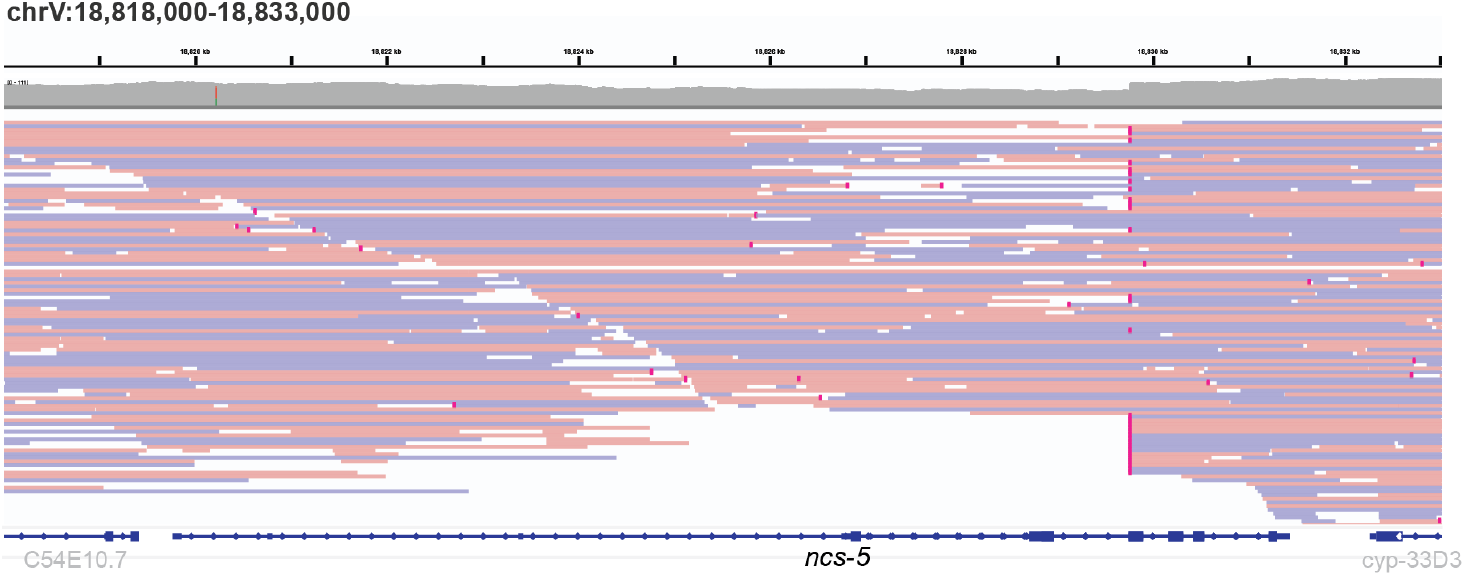
*maIs105[col-19p::gfp]* insertion candidate site on the right arm of Chromosome V. IGV genome browser screenshot of the identified strain-specific integration in VT3855 (*lin-46(ma467) maIs105*, Ilbay et al., 2021). Features are as in Figure 3. Similar to GR1395, soft-clipped reads indicate the insertion site of the transgene array. The array insertion origin of these reads is further supported by mappings of the clipped sequences to array components. The discontinuity in coverage and the presence of a subset of reads aligning through the integration site suggest a complex genomic rearrangement associated with the integration. a.Scatter plots comparing the log_2_ expression changes between the two biological replicates for the strains VT1367 vs. N2 (left panel) and GR1395 vs. N2 (right panel). Differentially expressed genes (see Methods) are colored in red. b.Karyo-type plot depicting the log_2_ expression changes as a function of genomic position. Each dot represents a gene. Red lines represent the array integration sites. Blue lines represent sites on chr. II also showing array components found in Nanopore sequencing of VT3855, to which VT1367 is the parental strain (see Methods).

### Two integrated multicopy transgene arrays cause gene expression changes and extensive promoter transcription

To begin exploring whether the presence of such large transgene arrays affects gene expression, we performed mRNA sequencing in early L1 stage VT1367 *maIs105[col-19p::gfp]* and GR1395 *mgIs49[mlt-10p::gfp::pest]* animals. (We selected early L1 as they are devoid of the high-amplitude oscillatory gene expression that occurs later in larval development and which can be a confounder in interpreting gene expression experiments involving single time point comparison (Aeschimann et al. 2017; Meeuse et al. 2020; Tsiairis and Großhans 2021; Bulteau and Francesconi 2022)). Each strain exhibited reproducible changes in expression relative to wild-type, when compared over two replicates each (Figure 5a). Using edgeR (Chen et al. 2025) with an FDR=0.05 and a minimum log_2_ fold change of 1, we found 59 down- and 76 upregulated genes in VT1367 *maIs105[col-19p::gfp]* and 273 down- and 199 upregulated genes in GR1395 *mgIs49[mlt-10p::gfp::pest]* (Figure 5a).

**Figure 5.**
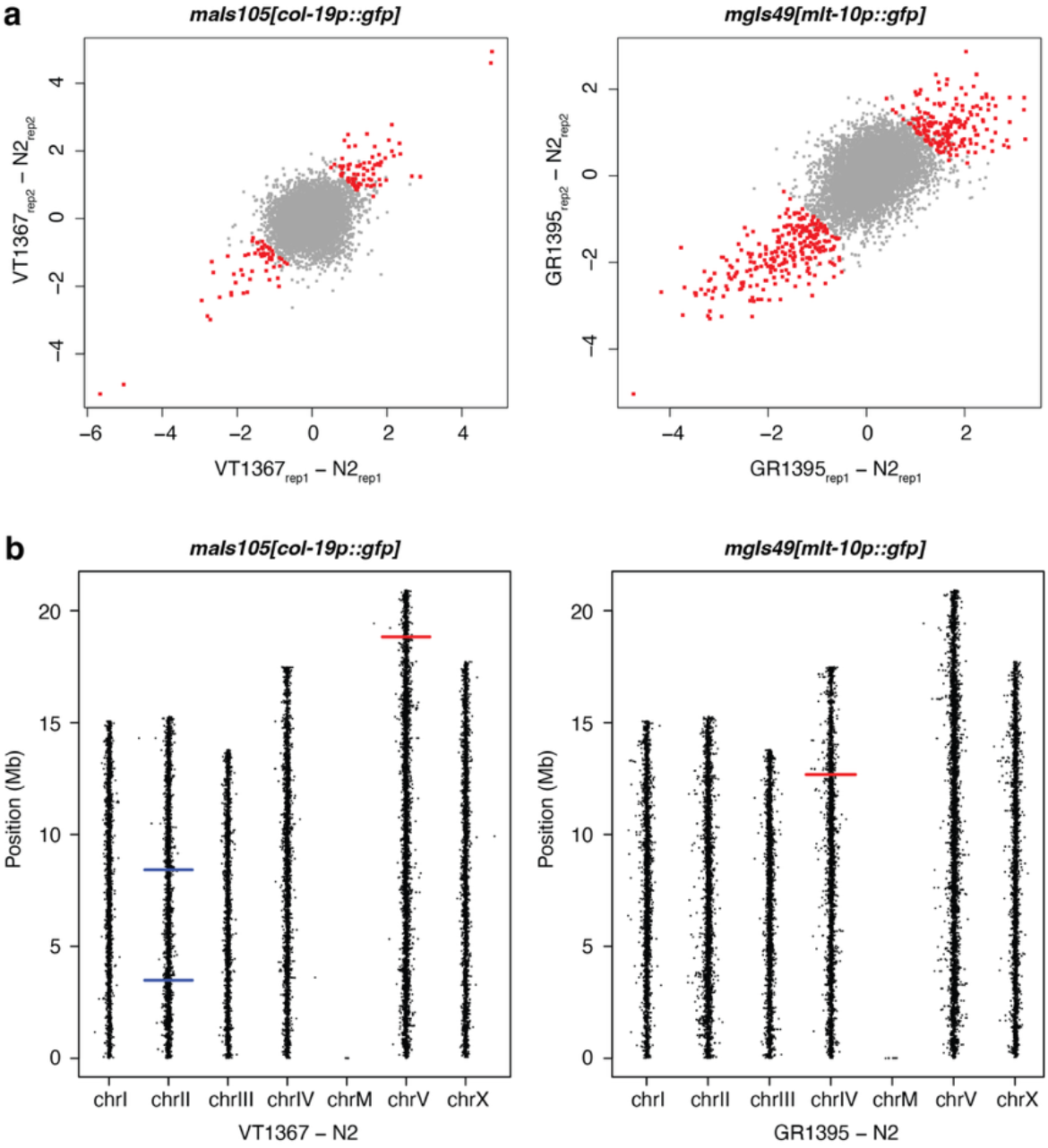
Integrated multicopy arrays cause gene expression changes.

We hypothesized that each array might preferentially perturb gene expression locally, i.e., close to its integration site, but saw no indication of such local expression perturbations (Figure 5b). However, we found evidence that there is extensive transcription of noncoding regions contained in each array: *mlt-10* (Frand et al. 2005) and *ttx-3* (Hobert et al. 1997) promoter in *mgIs49*-containing animals and *col-19* (Abrahante et al. 1998; Abbott et al. 2005) promoter in *maIs105*-containing animals, respectively (Figure 6).

**Figure 6.**
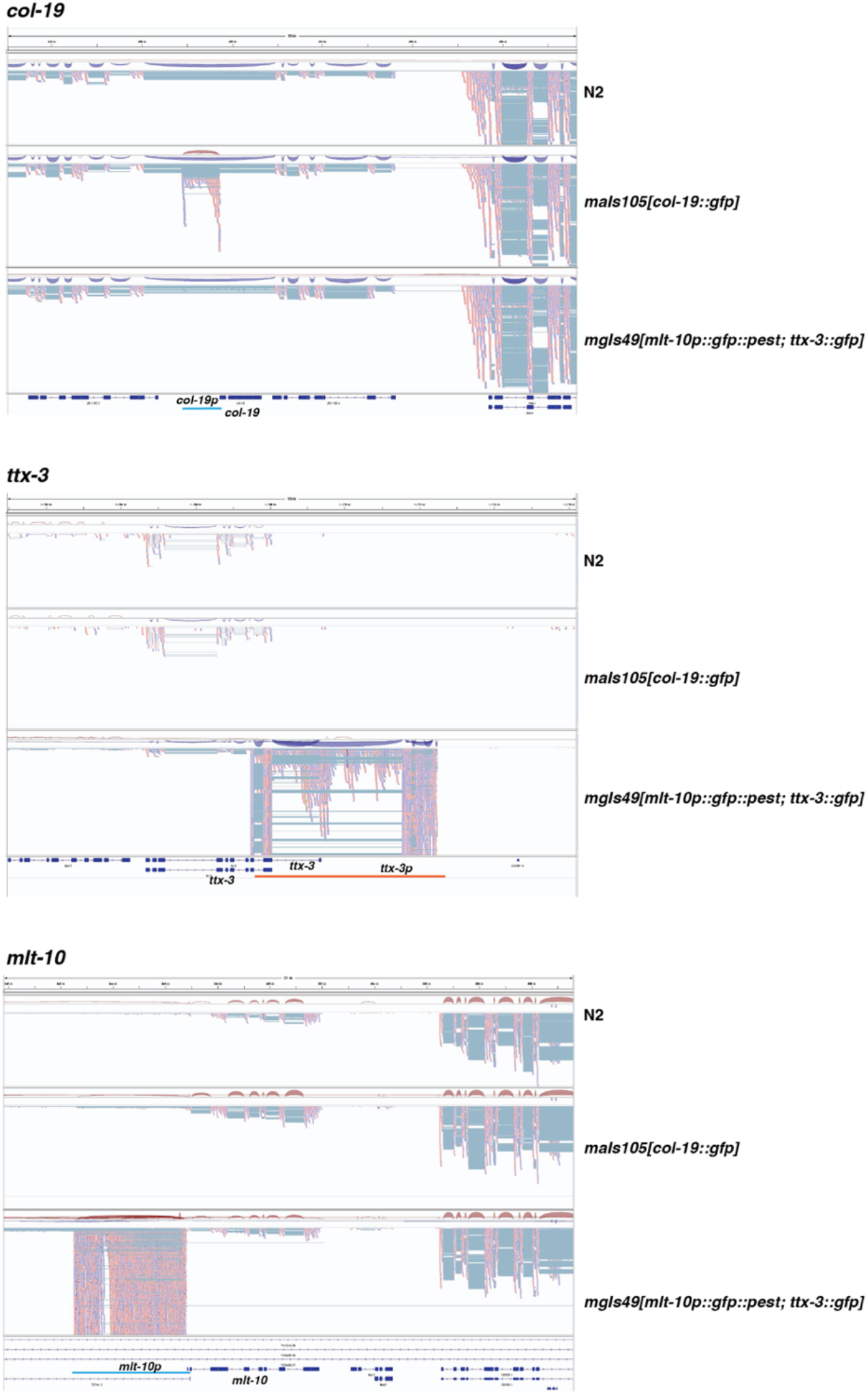
Transcripts covering transgene promoters accumulate in the transgene array-containing lines. Browser screenshots showing RNA-seq read alignments for the strains N2, VT1367 *maIs105[col-19p::gfp]* and GR1395 *mgIs49[mlt-10p::gfp::pest; ttx-3p::gfp]* at the *col-9, ttx-3* and *mlt-10* loci, respectively. Promoter regions present in the arrays are shown as orange or cyan lines, depending on gene orientation.

## Discussion

First established through pioneering DNA transformation experiments more than 30 years ago (Mello et al. 1991), multicopy transgene arrays have remained widely used tools in *C. elegans* research. Known shortcomings, such as proneness to silencing, especially in the germline, and overexpression artifacts have all but disqualified their use for gene expression, but they have remained popular as markers of cell fates and states. Our work provides a cautionary tale, showing that transgene arrays are problematic even for these more limited applications: Several arrays, both integrated and extrachromosomal, caused synthetic phenotypes with a *myrf-1* mutation.

Since the phenotypes were co-dependent on mutant *myrf-1*, one could consider the arrays as a means to sensitize the system. However, without knowing what exactly causes this sensitization, it seems difficult to interpret any phenotypes in a mechanistically meaningful way. This becomes even more obvious when such composite phenotypes are tested for modification, suppression or enhancement, through additional mutations – is the contribution of the array, the separate mutation, or the interaction between the two modified?

One hypothetical risk associated with transgene array integration is the disruption of host genes since integration occurs randomly. Our profiling data show that this is indeed what happened with the *mgIs49* array, which is integrated in, and disrupts, the *prmt-9* locus. However, although this may prove problematic in some contexts, it did not explain the *myrf-1* synthetic lethality. In principle, the effects of such insertional mutagenesis would also seem more controllable – since insertion occurs randomly, testing of a separate integrant in the genetic context of choice would readily rule out arrays causing specific defects. (In reality, however, the logistics of strain generation and testing make reporting of results from independent integrants a rare scenario.)

The fact that different transgene arrays, integrated as well as extrachromosomal, caused *myrf-1* synthetic lethality appears more concerning, pointing to a more general issue. Although the extent of synthetic lethality varied, the fact that it occurs independent of the component transgenes and integration status implicates more generic array features in the process, perhaps linked to their large size or repetitive nature.

We were indeed surprised by the large, megabase sizes that we generally observed for all eight integrated arrays that we had assessed. However, this number is in good agreement with other reports, examining distinct arrays: ONT Minion sequencing allowed assembly of an 11 Mb extrachromosomal array (Lin et al. 2021) while (Mouridi et al. 2022) estimated a simple integrated array size at 5.5 Mb. We note that the smallest array we identified, at 4.1 Mb, resulted from biolistic bombardment of a recombineering construct (Sarov et al. 2006). Although generally believed to be smaller, such arrays may be prone to introducing genomic rearrangements (Praitis et al. 2001; Tyson et al. 2018).

The risk of such rearrangements is likely a consequence of the DNA double strand repair rather than a specific consequence of biolistic transformation. Indeed, our preliminary analysis seeking to assign candidate integration sites for another seven sequenced strains, provided evidence for more than one possible integration site. For VT1367 *maIs105[col-19p::gfp]*, which we characterized in greater detail, ONT sequencing data analysis revealed three candidate integration sites, two on chromosome II and one on chromosome V. SNP mapping confirmed that GFP expression was linked to the right arm of chromosome V, but the coverage is discontinuous and a subset of reads align through the integration site in the *ncs-5* locus. We consider a complex genomic rearrangement as the most likely explanation of these observations.

Our data show that multiple arrays interact genetically with *myrf-1(mg412)*, but the ability to modulate mutant phenotypes does not appear limited to this allele. Thus, (Edelman et al. 2016) reported suppression of *lin-42(0)* mutant precocious alae formation by two out of three transgene arrays that they had tested. One of the suppressing arrays was *mgIs49*.

Finally, our mRNA sequencing data revealed substantial aberrant transcription in *mgIs49* and *maIs105*-carrying animals. While we do not know whether any of the mis-expressed genes contribute specifically to the *myrf-1* lethality phenotype, the accumulation of transcripts extending across the transgene promoters is striking. Whether these transcripts derive merely from the concatenation of truncated transgene copies remains to be determined, as does their functional relevance, but it seems quite possible that these aberrant transcripts could affect gene expression through anti-sense and other silencing mechanisms.

## Supporting information

Strain Table

## Acknowledgments

We thank Milou Meeuse for observing the loss of the extra molt phenotype upon outcrossing *myrf-1(mg412); mgIs49* animals. We thank the FMI Functional Genomics Platform for ONT sequencing and RNA sequencing our samples, Anca Neagu for the gift of the *mlt-10p* reporter plasmid, Kathrin Braun for advice on preparation of samples for RNA sequencing and luciferase assays, and Lucas Morales Moya for help with the luciferase analyzer.

## Funding

Some strains were provided by the CGC, which is funded by NIH Office of Research Infrastructure Programs (P40 OD010440). IK heads the Friedrich Miescher Institute for Biomedical Research (FMI) *C. elegans* facility, which is supported by FMI core funding. M.W.M.M. received support from a Boehringer Ingelheim Fonds PhD fellowship. This work is part of a project that has received funding from the Swiss National Science Foundation (#310030_207470, to H.G.) and from the European Research Council (ERC) under the European Union’s Horizon 2020 research and innovation programme (Grant agreement No. 741269, to H.G.). The FMI is core-funded by the Novartis Research Foundation

## Materials and methods

### General *C. elegans* methods

Nematodes were grown under standard conditions at 20° unless indicated otherwise. Mutant alleles were crossed into appropriate genetic backgrounds and, whenever feasible, a wild-type segregant from the same cross was used as a control.

Extrachromosomal arrays *bchEx30, bchEx31* and *bchEx32* are independent transgenic lines resulting from microinjecting the following mixture into adult gonads: *mlt-10p 4 kb::pest::gfp::h2b, unc-119(+)* at 10 ng/µl; *unc-122p::gfp* at 50 ng/µl; and 1 Kb Plus DNA Ladder (Invitrogen, 10787018) at 40 ng/µl.

### Scoring adult survival

Gravid adults were allowed to lay eggs for 2 hours and were subsequently removed from the plates, which were cultured at 25°C. 24 hours later, larvae were counted and transferred to fresh plates. Surviving adults were counted at 96 hours.

### Genome editing by CRISPR/Cas9

Genome editing was performed as described by (Ghanta and Mello 2020).

*myrf-1(bch65)* codes for a T364A missense mutation and was created in the N2 genetic background. It also includes silent mutations R370R, L371L, H372H. *myrf-1(bch71), myrf-1(bch72) and myrf-1(bch73)* are identical alleles that revert the *myrf-1(mg412)* mutation to the wild-type sequence. They were created in the background of the strain IFM363 *myrf-1(mg412) II; mgIs49 [mlt-10p::gfp::pest + ttx-3::gfp] IV; xeSi311 [eft-3p::luc::gfp::unc-54 3’UTR, unc-119(+)] V. prmt-9(bch79), prmt-9(bch80)* and *prmt-9(bch81)* are identical *prmt-9* truncation alleles, created in the background of the strain IFM332 *myrf-1(bch65[T364A])*.

### Single-copy transgene integration by MosSCI

Transgenic worms expressing the *mlt-10p 4 kb::pest::gfp::h2b, unc-119(+)* construct were obtained by single-copy integration into the site *oxTi444* on chromosome III (Frøkjær-Jensen et al. 2012) to yield *bchSi134 [mlt-10p::pest::gfp::h2b]* and *bchSi135 [mlt-10p::pest::gfp::h2b]*.

### Luciferase assays

Luciferase assays were performed as described (Meeuse et al. 2020). Briefly, gravid adults were treated with a bleaching solution to extract eggs. Single embryos were transferred into a well of a white, flat-bottom, 384-well plate (Berthold Technologies, 32505) by pipetting. Animals developed in 90 mL S-Basal medium containing *E. coli* OP50 (OD600 = 0.9) and 100 mM Firefly D-Luciferin (p.j.k., 102111). Plates were sealed with Breathe Easier sealing membrane (Diversified Biotech, BERM-2000). Luminescence was measured using a luminometer (Berthold Technologies, Centro XS3 LB 960) every 10 min for 0.5 s for 72 or 120 hours; the animals experience a temperature of 22°C. An automated algorithm was used to detect the hatch and the molts from the luminescence data.

### GFP imaging

Fluorescent and Differential Interference Contrast (DIC) images were acquired using a Zeiss Axio Observer Z1 microscope with AxioVision software and Zen 2 (Blue Edition). Region selection and image processing was performed using Fiji.

### Oxford Nanopore DNA sequencing

Mixed-stage animals were grown on a 10 cm NGM plate and washed three times with M9 buffer, pelleted and frozen in a dry ice-ethanol bath for 10 minutes. The DNA was extracted using the QIAGEN Puregene Cell Kit (158043) and Proteinase K (19131) as described in the **Supplemental Protocol**.

### DNA was eluted in 200 µl water per sample

4µg of genomic DNA was fragmented to a target size of 15kb using G-Tubes (Covaris). Library preparation was performed using the Native barcoding kit v14 (SQK-NBD114.24) from Oxford Nanopore Technologies, following the manufacturer recommendations. Pooled libraries were sequenced on a PromethION flowcell (R10.4.1) using a P2 solo device.

### Processing of Oxford Nanopore DNA sequencing data

Raw POD5 signal files were basecalled using Dorado (version 0.8.1) from Oxford Nanopore Technologies with the R10.4.1 high-accuracy model (dna_r10.4.1_e8.2_400bps_hac@v5.0.0). Basecalling was performed on an NVIDIA A40 GPU. Basecalled reads were demultiplexed using dorado demux (kit SQK-NBD114-24) with barcode trimming enabled, yielding per-barcode read sets corresponding to the analyzed strains. Demultiplexed long reads were subsequently aligned to the C. *elegans* genome (ce11) using minimap2 (version 2.24) with the map-ont preset. Alignments were converted to BAM, sorted by coordinate, and low-confidence mappings were removed (minimum Q-score threshold of 8). These per-strain BAM files were used for downstream insertion-site discovery and array size estimation.

### Identification of candidate insertion sites from ONT reads

To identify genomic insertion sites of large integrated transgenic arrays, aligned ONT reads from each strain were analyzed for breakpoint-like alignment signatures. We first extracted reads that showed at least 500 bp of terminal soft clipping, as expected for junction-spanning reads containing genomic sequence contiguous with non-reference array DNA. For each read, the genomic breakpoint coordinate was defined from the alignment boundary, taking strand orientation into account, and the number of reads supporting each chromosome-position pair was recorded and only candidate breakpoint sites with a minimum support of 5 reads were retained. To remove recurrent background signals, breakpoint coordinates detected in all other strains were pooled and subtracted from those of the target strain, retaining only strain-specific sites. For each remaining candidate, we then determined how many reads overlapping the genomic breakpoint also aligned to reference sequences representing known array components. Candidate insertion sites were defined as strain-specific breakpoint hotspots supported by multiple soft-clipped reads, including reads with matches to array-derived sequence. Finally, insertion sites were confirmed by visual inspection in a genome browser.

### Estimation of total array sizes from ONT reads mapping to array components

Total array sizes (over all possible insertions) were estimated from the relative number of long reads mapping to array components versus those mapping to the genome. For each sample, ONT reads were aligned to reference sequences representing known construct components using minimap2 (Li 2021) and the map-ont preset. Unmapped, secondary, and supplementary alignments were excluded from the construct-component BAM, and the corresponding genome-aligned BAM was used for normalization. Each read was counted only once per BAM file and its length was summed to obtain the total number of mapped base pairs for construct-component and genome alignments. Genome depth was then estimated as mapped genomic base pairs divided by the haploid C. *elegans* genome size. Finally, total array size per diploid genome was calculated by normalizing construct-mapped base pairs by this genome depth.

### RNA sequencing

N2, GR1935 *mgIs49 [mlt-10p::gfp::pest; ttx-3::gfp]* and VT1367 *maIs105 [col-19::gfp]* L1 animals through starvation by hatching eggs into M9 buffer overnight. Following plating on OP50-seeded plates, animals were allowed to develop for 4 hours at 25°C. The animals were washed three times and lysed in Trizol by repeated freeze-thawing cycles. RNA was extracted using the Direct-zol RNA Microprep kit (Zymo, R2062). mRNA-seq libraries were generated using the Illumina Stranded mRNA Prep solution, according to the manufacturer’s protocol. Libraries were sequenced on a NovaSeq6000 flowcell, paired-end 2x56cycles. Demultiplexing was performed using bcl2fastq2.

### Processing of the RNA-seq data

RNA-seq data was mapped to the *C. elegans* genome (ce11) using the align (qAlign) function from the QuasR (version 1.34.0) package in R with splicedAlignement = TRUE, using the aligner HISAT2 (version 1.10.0) including an exon-exon junction database. Gene expression levels were quantified using the qCount() function. For annotations, coding transcript info from WormBase WS270 was used. We further compensated for differences in library sizes by scaling each library to the average library size, and log2-transformed the data using a pseudocount of 8 (log_2_(x+8)). Given that fragments of the genes of *mlt-10, ttx-3* and *col-19* occur in the arrays, we removed the affected exons from those genes to only count reads that originated from the endogenous genes as opposed to the array. To determine those affected exons, we inspected the read alignments at those loci and removed the exons at the 5’ end that overlapped sections with extensive read amplification. Differentially expressed genes between GR1395 *mgIs49 [mlt-10p::gfp::pest; ttx-3::gfp]* and N2 as well as VT1367 *maIs105 [col-19::gfp]* and N2, respectively, were identified using the R package edgeR (version 4.8.2) (Chen et al. 2025). Genes with low expression were filtered out using filterByExpr. Quasi-likelihood F-tests were performed to assess differential expression, and genes with a false discovery rate (FDR) < 0.05 and minimum log_2_ fold change of 1 were considered significantly differentially expressed.

### Genetic mapping of *maIs105*

VT1367 *maIs105 [col-19::gfp]* hermaphrodites were mated with CB4856 Hawaiian males. Two kinds of F_2_ animals were analyzed: those homozygous for the array and those homozygous for its absence. SNP analysis using the SNPs pkP5076 (V, −17), pkP5097 (V, 1), *** (V,6), uCE5-2609 (V, 13) and snp_Y17D7B[3] (V, 18) (Davis et al. 2005) of 72 animals placed *maIs105* on the right arm of chromosome V (distal to snp_Y17D7B [3]).

## Data availability

The sequencing data produced in this study are available at NCBI’s Gene Expression Omnibus (Edgar et al. 2002) under SuperSeries accession number GSE329497 (https://www.ncbi.nlm.nih.gov/geo/query/acc.cgi?acc=GSE329497).

## Supplemental Protocol

DNA extraction from frozen worm pellets for Oxford Nanopore Sequencing, based on https://nanoporetech.com/document/extraction-method/c-elegans-dna

QIAGEN Puregene Cell Kit (158043) and Proteinase K (19131)

1. Add 1.5 ml of Cell Lysis Solution to the 15 ml Falcon tube containing the frozen worm pellet, and allow the pellet to thaw in the lysis buffer.
2. Add 7.5 µl of Proteinase K and resuspend the pellet by pipetting with a 1 ml wide-bore tip.
3. Incubate the resuspended worms at 50°C for 1 hour; During this incubation, gently invert the tube 3 times every 30 minutes.
4. Repeat steps 1-3 once more.
5. Add 15 µl of RNase A and mix by inverting the tube.
6. Incubate the tube at 37°C for 30 minutes.
7. Place the tube on ice for 2 minutes.
8. Add 1 ml of Protein Precipitation Solution and pulse-vortex three times for 5 seconds.
9. Centrifuge at 2000 x g for 10 minutes.
10. Add 3 ml of isopropanol to a fresh 15 ml Falcon tube.
11. Pour the supernatant from step 9 into the Falcon tube with isopropanol. Discard the pellet.
12. Gently invert the tube 50 times.
13. Centrifuge at 2000 x g for 5 minutes.
14. Discard the supernatant and add 3 ml of 70% ice-cold ethanol to the pellet. Gently invert the tube several times to mix.
15. Centrifuge at 2000 x g for 2 minutes.
16. Discard the supernatant and remove as much ethanol as possible using sterile paper wipes.
17. Add 200 µl water, vortex for 5s. Incubate at 65 °C for an hour. Incubate overnight with gentle shaking.

**Supplementary Table 1:**
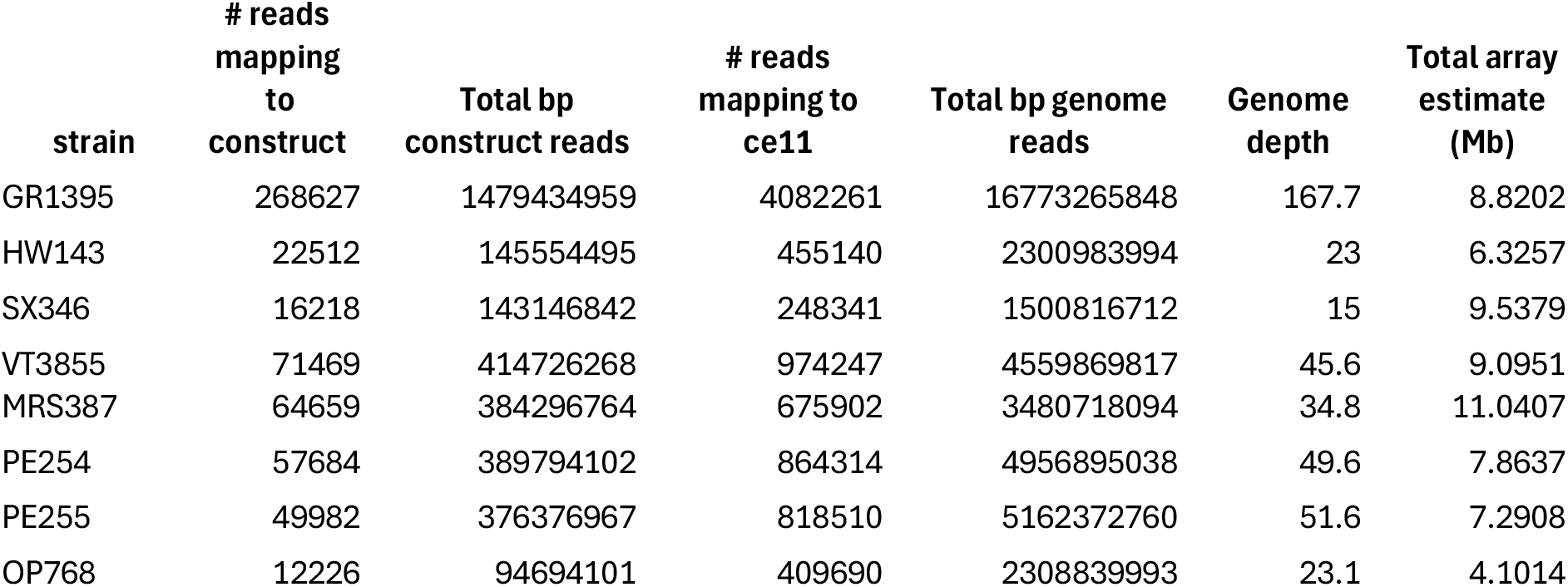
Array size estimation.

**Supplementary Table 2:** Candidate integration sites.

| Strain | Integration<br>Locus | Chromosome | Coordinate | Genomic<br>Support | Array<br>Support | Breakpoint<br>Support |
| --- | --- | --- | --- | --- | --- | --- |
| OP768 | 1/2 | chrV | 1986898 | 54 | 12 | 10 |
| OP768 | 2/2 | chrV | 2344706 | 43 | 12 | 5 |
| PE255 | 1/2 | chrX | 4257408 | 27 | 10 | 12 |
| PE255 | 2/2 | chrX | 17200018 | 81 | 22 | 11 |
| PE254 | 1/1 | chrV | 20165630 | 74 | 34 | 17 |
| MRS387 | 1/1 | chrX | 14209166 | 19 | 14 | 14 |
| MRS387 | 1/1 | chrX | 14209174 | 21 | 20 | 8 |
| VT3855 | 1/3 | chrV | 18829757 | 91 | 28 | 10 |
| VT3855 | 2/3 | chrII | 3483183 | 59 | 20 | 15 |
| VT3855 | 3/3 | chrII | 8420157 | 30 | 29 | 12 |
| SX346 | 1/2 | chrII | 14770046 | 10 | 9 | 6 |
| SX346 | 2/2 | chrX | 5472798 | 53 | 31 | 23 |
| SX346 | 2/2 | chrX | 5478852 | 75 | 14 | 5 |
| SX346 | 2/2 | chrX | 5479078 | 76 | 14 | 5 |
| SX346 | 2/2 | chrX | 5483955 | 48 | 25 | 21 |
| HW143 | 1/2 | chrV | 14013649 | 48 | 31 | 7 |
| HW143 | 2/2 | chrV | 7068894 | 47 | 14 | 7 |
| GR1395 | 1/1 | chrIV | 12677932 | 81 | 79 | 28 |
| GR1395 | 1/1 | chrIV | 12679597 | 101 | 16 | 48 |

## Notes

### Competing Interest Statement

The authors have declared no competing interest.

